# Physico-chemical principles of HDL-small RNA binding interactions

**DOI:** 10.1101/2021.12.17.473011

**Authors:** Danielle L. Michell, Ryan M. Allen, Ashley B. Cavnar, Danielle M. Contreras, Minzhi Yu, Elizabeth M. Semler, Marisol A. Ramirez, Wanying Zhu, Linda May-Zhang, Chase A. Raby, Mark Castleberry, Anca Ifrim, John Jeffrey Carr, James G. Terry, Anna Schwendeman, Sean S. Davies, Quanhu Sheng, MacRae F. Linton, Kasey C. Vickers

**Author notes:** Correspondence to: Danielle L. Michell, PhD, 2220 Pierce Ave. 312 Preston Research Building, Nashville, TN. 37232 USA, or, Kasey C. Vickers, PhD, 2220 Pierce Ave. 312 Preston Research Building, Nashville, TN. 37232 USA.

## Abstract

Extracellular small RNAs (sRNA) are abundant in many biofluids, but little is known about their mechanisms of transport and stability in RNase-rich environments. We previously reported that high-density lipoproteins (HDL) of mice were enriched with multiple classes of sRNA derived from the endogenous transcriptome, but also exogenous organisms. Here, we show that human HDL transports tRNA-derived sRNAs (tDRs) from host and non-host species which were found to be altered in human atherosclerosis. We hypothesized that HDL binds to tDRs through apolipoprotein A-I (apoA-I) and these interactions are conferred by RNA-specific features. We tested this using microscale thermophoresis and electrophoretic mobility shift assays and found that HDL bind tDRs and other single-stranded sRNAs with strong affinity, but not doublestranded RNA or DNA. Natural and synthetic RNA modifications influenced tDR binding to HDL. Reconstituted HDL bound tDRs only in the presence of apoA-I and purified apoA-I alone was sufficient for binding sRNA. Conversely, phosphatidylcholine vesicles did not bind tDRs. In summary, HDL preferentially binds to single-stranded sRNAs likely through non-ionic interactions with apoA-I. These studies highlight binding properties that likely enable extracellular RNA communication and provide a foundation for future studies to manipulate HDL-sRNA for therapeutic approaches to prevent or treat disease.

## INTRODUCTION

High-density lipoproteins (HDL) have many beneficial properties which antagonize inflammation, infection, metabolic dysfunction, and disease[1, 2]. HDL is the smallest class of lipoproteins (7-12 nm in diameter) and the HDL pool is composed of a compendium of sub-species with different sizes, shapes, and functions[1]. Apolipoprotein A-I (apoA-I), the primary structural and functional protein of HDL, is secreted from hepatocytes and intestinal enterocytes[3]. Upon secretion, apoA-I acquires phospholipids and free cholesterol from cells to form nascent pre-β discoidal HDL particles[4, 5]. Further acceptance and esterification of free cholesterol cargo results in the formation of mature spherical particles containing a hydrophobic core with a phospholipid monolayer shell. HDL particles are dynamic and continuously load and unload cargo with interacting cells and other lipoproteins. The HDL pool contains >215 different proteins; however, apoA-I accounts for approximately 70% of HDL total protein mass[6, 7]. In addition to proteins and diverse bioactive lipids, HDL also transport many classes of small non-coding RNAs (sRNAs)[8, 9] in both plasma and biofluids (e.g. lymph) and have been shown to transport and deliver functionally active microRNAs (miRNA) to recipient cells where they participate in gene regulation networks [10–13]. We reported that HDL’s anti-inflammatory capacity in endothelial cells was found to be mediated, in part, by HDL delivery of miR-223-3p to recipient human coronary artery endothelial cells and silencing of intercellular adhesion molecule 1[10]. The HDL-miRNA profile is significantly altered in multiple diseases, including familial hypercholesterolemia, atherosclerosis, uremia, diabetes, and obesity[9, 14–17]. HDL-miRNAs have also been reported to be altered by diet and specific dietary components[18–20]. To date, most investigation of extracellular sRNAs on lipoproteins has been limited to miRNAs; however, miRNAs only represent a small fraction of sRNAs on HDL and other classes of sRNA warrant investigation [8]. Recently, we profiled HDL-sRNAs in mice using high-throughput sRNA sequencing and many classes of host and non-host sRNAs were identified on circulating HDL[8]. Results showed that HDL also transport tRNA-derived sRNAs (tDRs) and rRNA-derived sRNAs (rDR)[8]. Furthermore, we found that mouse HDL are highly-enriched with non-host microbial sRNAs which likely originate from bacteria and fungi in the microbiome, environment, and/or diet.

HDL-sRNAs are likely single-stranded and generally <60 nts in length[8, 9]. HDL have been found to accept sRNAs from multiple cell types, including miR-223-3p from macrophages and neutrophils, and miR-375-3p from pancreatic beta cells[9, 21, 22]. Nevertheless, the mechanism by which HDL acquire sRNAs from donor cells and the underlying biochemistry of HDL-sRNA binding have remained elusive. Studies suggest that oligonucleotides can associate with zwitterionic lipids, e.g. phosphatidylcholine (PC)[23–26], but it is currently unknown if natural PC within HDL’s shell mediate it’s transport of sRNA. Results from HDL delipidation studies suggest that miRNAs may bind to HDL proteins as opposed to phospholipids[14, 27]; however, to date, a candidate protein on HDL with RNA binding capacity has not been identified. Here, we tested the hypothesis that apoA-I is a novel RNA binding protein for single-stranded sRNAs, specifically tDRs. Microscale thermophoresis (MST) and electrophoretic mobility shift assay (EMSA) data suggest that apoA-I is a novel ribonucleoprotein that likely confers the ability of HDL to bind RNA, rather than the lipid cargo of the particle. MST approaches were used to define the impact of base modifications and the 2’ hydroxyl group on HDL-sRNA binding. These results support a mechanism by which HDL binds sRNAs, including tDRs, and reveal the physicochemical properties that influence HDL-sRNA binding.

## MATERIALS AND METHODS

### Reagents

Synthesized oligonucleotides used in this study are listed in **Table S2**.

### Clinical Assays

Plasma was collected from whole venous blood (K_2_EDTA) obtained from participant donors following informed written consent. Experiments were performed under institutional review board approved protocols from Vanderbilt University Medical Center (IRB#170046 and #101615). Participants underwent non-contrast coronary computed tomography (CT) examination to identify the presence of calcified coronary atheroma and measure the coronary artery calcium (CAC) score[28, 29]. Women who were pregnant or potentially pregnant were excluded from the CT examination. Image acquisition was performed using a 64-slice multi-detector CT scanner (Brilliance 64, Philips Healthcare, Cleveland, OH, USA). Technical parameters included: 120 KVp, 150 mAs, and ECG-gating of image acquisition in late diastole. CAC was measured on 3.0 mm thick slices using the Food and Drug Administration–approved calcium scoring software (Philips IntelliSpace Portal, Philips Healthcare, Cleveland, OH, USA) and reported as the Agatston score[30] for minimum lesion volume of 0.5 mm3 and an attenuation threshold of ≥130 Hounsfield units. Subject characteristics are presented in **Table S1**. Plasma lipoprotein cholesterol and plasma triglyceride (TG) levels were quantified using an Ace Axcel clinical chemistry system (Alfa Wassermann).

### Lipoprotein Isolation

Density-gradient ultracentrifugation (DGUC) was completed, as previously reported[31]. Native very low-density lipoproteins (VLDL, 1.006-1.018g/L), low-density lipoproteins (LDL) (1.019-1.062g/L), and HDL (1.063-1.021g/L) were isolated by sequential DGUC using an Optima XPN-80 Ultracentrifuge with SW32Ti or SW41Ti rotors (Beckman-Coulter). Lipoproteins were dialyzed in 1X PBS (4L) for at least 4 bucket changes and concentrated with 3 kDa m.w. cutoff filters (Millipore). Isolated human DGUC-HDL, cell culture media, or plasma samples were injected into an AKTA Pure fast-protein liquid chromatography system (FPLC, Cytiva) with three tandem Superdex-200 Increase size-exclusion chromatography (SEC) columns (Cytiva) and collected in 1.5mL fractions in running buffer (10mM Tris-HCl, 0.15M NaCl, 0.2% NaN3), as previously described(21). SEC fraction fluorescence was measured with microplate fluorometer (SynergyMx, Biotek). DGUC-HDL or individual SEC fractions were assessed for total protein (Pierce BCA, ThermoFisher), total cholesterol (TC), PC, and TG (Pointe Scientific) by colorimetric assays.

### HDL Delipidation

DGUC-HDL were delipidated as previously described[10]. Briefly, 6mL of ice cold methanol:chloroform (1:1) was added to 1mL of lyophilized HDL, mixed, and incubated on ice for 30min. Chilled methanol (4mL) was added to the mixture, centrifuged (3,000 RPM) for 10min, removed supernatants, and repeated until pellet turned white. Chilled methanol (10mL) was re-added, mixed, and recentrifuged. The pellet was dried under nitrogen, resolubilized with 6M guanidine HCl, and dialyzed in ammonium bicarbonate (10mM) overnight. ApoA-I was isolated from delipidated HDL with 6M Urea by anion exchange chromatography with Q-Sepharose Fast Flow column coupled with a Q-Sepharose High-Performance column on an AKTA Pure FPLC System (Cytiva). ApoA-I fractions were confirmed using Coomassie-based Aquastain (BulldogBio) on an SDS-PAGE gel and were dialyzed in ammonium bicarbonate (10mM).

### HDL Reconstitution (rHDL)

rHDL was prepared with apoA-I and L-α-PC (Avanti Polar Lipids) at 100:1 (PC:Protein). Lyophilized PC was resuspended in PBS (80μL:1mg PC) and vortexed. 10% sodium deoxycholate was added (1/5 total volume), vortexed, and incubated at 37°C for 30min in water bath. Purified apoA-I was added to the reaction and was incubated at 37°C for 1h. Samples were dialyzed and rHDL were assessed by SEC. As a control, small unilamellar vesicles (SUV) were prepared using evaporated Egg PC (Avanti Polar Lipids) and HEPES Buffered Solution (20mM HEPES, 150mM NaCl) and incubated for 1h RT. Samples were vortexed vigorously, sonicated, and prepared SUVs were analyzed by SEC and stored at 4°C.

### Preparation of synthetic HDL (sHDLs)

22A peptide (PVLDLFRELLNELLEALKQKLK) was synthesized by Genscript. 1-Palmitoyl-2-oleoyl-sn-glycero-3-phosphocholine (POPC), 1,2-dimyristoyl-sn-glycero-3-phosphocholine (DMPC), 1,2-dipalmitoyl-sn-glycero-3-phosphocholine (DPPC), 1,2-distearoyl-sn-glycero-3-phosphocholine (DSPC), 1,2-dipalmitoyl-sn-glycero-3-phosphorylglycerol sodium salt (DPPG-Na), and 1,2-dimyristoyl-sn-glycero-3-phosphoethanol-amine (DMPE) were purchased from NOF America Corporation. 22A peptide and phospholipids were dissolved in acetic acid and mixed at a peptide/lipid mass ratio of 1:2 and lyophilized overnight. The lyophilized powder was rehydrated with phosphate-buffered saline (PBS; pH 7.4). The solution was then cycled 3 times between 50 °C (10min) and room temperature (10min) to allow the formation of sHDL complexes. The hydrodynamic size of the resulting sHDLs was analyzed by dynamic light scattering (DLS) using Zetasizer Nano ZSP (Malvern Instruments). The purity of sHDLs was analyzed by gel permeation chromatography (GPC) using a Tosoh TSK gel G3000SWxl column (Tosoh Bioscience) with UV detection at 220nm.

### HDL dicarbonyl modifications

To modify HDL with reactive dicarbonyls modifications, DGUC-HDL were reacted with 1 molar equivalence (eq) isolevuglandin (IsoLG) to HDL-apoA-I protein levels, 1 molar eq of 4-oxo-2-nonenal (ONE), or control vehicle dimethylsulfoxide (DMSO). The reaction was completed overnight at 37°C.

### Microscale thermophoresis (MST)

Synthesized AF647-labelled oligoribonucleotides (100nM, IDT, **Table S2**), denatured at 90°C for 2min and then cooled, were incubated (1:1) with 16 serial dilutions of ligand (e.g. HDL) in PBS at RT for 5min protected from light. sRNA-ligand solutions were loaded into standard-treated glass capillaries (NanoTemper Technologies) and tested using a Monolith NT.115 instrument (NanoTemper Technologies). Other buffers also include PBS with potassium acetate (Research Products International) and DPBS with CaCl_2_ and MgCl_2_ (Corning). For all MST experiments, 20% excitation power and 40% MST power were used. Each capillary was scanned to assess fluorescence homogeneity and fluorescence adsorption to ensure adequate sample quality, and MST measurements were then performed using the MO. Control software (v1.6, NanoTemper Technologies). Measurements were performed in triplicate and data analyses were performed on the MO.Affinity Analysis software (v2.3, NanoTemper Technologies) with representative binding curves presented. The dissociation constant (K_d_) was determined by the MO. Affinity Analysis software.

### Electrophoretic Mobility Shift Assays (EMSA)

EMSA were performed as previously described[32]. Briefly, AF546-tDR-GlyGCC-30 (1μM) was incubated with HDL (serial dilutions) and crosslinked (UV 254nm) for 15min on ice. Cross-linked samples were separated by native gel electrophoresis – NativePAGE 4-16% gel (ThermoFisher) with 4X NativePAGE sample buffer (ThermoFisher) and 0.5X TBE running buffer. Fluorescence signal (representing migration of tDR-GlyGCC-30) in the gel was visualized with Green Epi Illumination and 602/50 emission filter using a Digital ChemiDoc MP (BioRad) and protein-staining was completed by Coomassie-based Aquastain (Bulldog Bio). Sample protein masses were assessed with Amersham high molecular weight protein marker mix (GE Healthcare) with proteins porcine thyroglobulin (R_s_=17.0nm), equine spleen ferritin (R_s_=12.2nm), bovine heart lactate dehydrogenase (R_s_=8.2nm) and bovine serum albumin (R_s_=7.1nm) [33].

### Animals

Wild type C57BL/6 mice from The Jackson Laboratory were used under approved Vanderbilt Institutional Animal Care and Usage Committee protocols. Mice were housed in a 12h light/dark cycle with unrestricted access to standard chow (NIH-31) and water. Male 8-10 week-old mice were used in this study.

### Tissue Culture

Human peripheral blood mononuclear cells (PBMCs) were isolated by Ficoll-gradient centrifugation, washed, and counted. PBMCs were incubated with anti-human CD14 MicroBeads (Miltenyi Biotec) and monocytes were separated on a LS magnetic column on a QuadroMACS separator (Miltenyi Biotec). Cells were differentiated with human GM-CSF (50ng/mL, Sino Biological) in RPMI 1640 medium with L-glutamine (300mg/L, ThermoFisher), supplemented with 10% heat-inactivated fetal-bovine serum (FBS, ThermoFisher) and 1% penicillin/streptomycin (Gibco) for 7d followed by 24h of native HDL (nHDL, 1mg/mL) treatments in FBS-free media. Bone marrow cells were isolated from wild-type C57BL/6 mouse femurs and differentiated to bone marrow-derived macrophages (BMDM) using murine GM-CSF (50ng/mL, Tonbo Biosciences) in DMEM (ThermoFisher) supplemented with sodium pyruvate (110mg/L), L-glutamine (584mg/L), 10% heat-inactivated FBS (ThermoFisher), and 1% penicillin/streptomycin (Gibco) for 7d. BMDMs were transiently-transfected with tDR-GlyGCC-30-AF647 (1μg/mL) complexed to DOTAP transfection reagent (Roche) for 24h in serum-free media followed by nHDL (1mg/mL) treatments in FBS-free media for 24h.

### Protein Assays

For western blotting, samples were denatured in 4X sample buffer (Li-Cor Biosciences) with 10% β-mercaptoethanol at 70°C for 10min and separated by gel electrophoresis (NuPAGE 4-12% Bis-Tris gel with 1X MES running buffer, ThermoFisher). Gels were transferred to nitrocellulose membranes (ThermoFisher), blocked in Odyssey blocking buffer (TBS, Li-Cor Biosciences) for 30min, and incubated overnight with rocking at 4°C with anti-human apoA-I primary antibody (mouse monoclonal, 1:8000, Meridian Life Sciences). Membranes were washed 3X for 10min with 1X TBS-T (0.1% Tween 20) and incubated with anti-mouse secondary antibody conjugated to IRDye 800CW (Li-Cor Biosciences) for 1h. Membranes were washed (3 x 10min) and imaged with the LI-COR Odyssey Infrared Imaging system and quantified using Image Studio Lite and ImageJ software suites. Precision Plus Kaleidoscope Pre-Stained Protein Standards were used for ladder (BioRad). For enzyme-linked immunoassays (ELISA), human Apolioprotein A1 ELISA^PRO^ kits (Mabtech) were used to quantify apoA-I levels (as normalized to total protein) on HDL from CAC and healthy donors.

### RNA Assays

DGUC-HDL isolated from CAC and healthy donors were assayed for total RNA content using SYTO RNASelect stain (ThermoFisher). DGUC-HDL was incubated with 50μM of SYTO stain, rocked at 37°C at 225 RPM for 25min, and quantified by fluorometry (490/530nm). HDL total RNA levels were determined using a standard curve of single-stranded oligoribonucleotide (**Table S2**), and total RNA levels were normalized to total protein levels. Total RNA were isolated from equivalent amounts of HDL total protein using the miRNeasy Mini Kit (Qiagen). For sRNA analyses, RNA were reverse transcribed using the miRCURY LNA RT kit (Qiagen) and qPCR was performed using custom primers (**Table S2**) and PowerUp SYBR Green 2X Master Mix (ThermoFisher) on QuantStudio6 or 12k Real-Time PCR instruments (ThermoFisher). Relative quantitative values (RQV) were determined using an arbitrary Ct of 32 (RQV = 2^−ΔCTArb^).

### sRNA sequencing

HDL-sRNA libraries were generated using the NEXTFlex Small RNA-Seq Kit v3 with 1:4 adapter dilution and 22 PCR cycles (PerkinElmer). Following PCR amplification, individual libraries were size-selected (136-200bp) using Pippin-Prep (Sage Science) by 3% agarose gel cassette with Marker P dye (Sage Science). Libraries (cDNA) were quantified by Qubit (High-sensitivity DNA assay kit, ThermoFisher) and checked for quality by bioanalyzer (High-sensitivity DNA chips, 2100 Bioanalyzer, Agilent) prior to multiplexing using equal molar concentrations, and concentrated (DNA Clean and Concentrator kit, Zymo) for multiplex sequencing on the NextSeq500 (Illumina) by the Vanderbilt Technologies for Advanced Genomics (VANTAGE) core (Vanderbilt University Medical Center, Nashville, TN). Samples were demultiplexed and analyzed using the TIGER pipeline[8]. Data analyses were completed using the TIGER sRNA pipeline, as previously described.

### Statistics

To determine differential expression (abundance) of HDL-sRNAs by sRNA-seq, DESeq2 analyses were applied as part of the TIGER pipeline[34]. For comparisons between two independent groups, unpaired two-tailed t-tests were used. For correlation analyses, Pearson correlations were used. A p-value <0.05 was considered significant. GraphPad Prism 9 (GraphPad Software) was used to generate graphs. Affinity Designer was used to assemble the images and generate the figures.

## RESULTS

### Human HDL transports diverse sRNA classes (Fig.1)

The determination of overall total RNA content on human HDL particles has previously been challenging due to technical limitations that require RNA isolation and downstream quantitation of individual candidate sRNAs of prior knowledge. To overcome these barriers, total RNA levels were quantified in intact human HDL with SYTO RNASelect, a fluorescence-based membrane-penetrating RNA dye that does not require RNA isolation[35]. HDL were isolated from healthy controls (Control, mean=0, N=46) and subjects with indications of atherosclerosis (CAC, mean=61, n=21) (**Fig.1A, Table S1**). Based on an oligoribonucleotide standard curve, HDL particles were determined to transport about 128μg of total RNA per mg HDL total protein for healthy control subjects (n=16) and 125μg total RNA per mg HDL total protein for CAC subjects (n=11) (**Fig.1B**). In both control and CAC subjects, DGUC-HDL total cholesterol levels were significantly correlated to HDL-RNA (**Fig.1C**). HDL-apoA-I protein levels were also found to be significantly correlated to HDL-RNA levels in control subjects, but not CAC subjects (**Figs.1D, S1A, B**).

**Figure 1.**
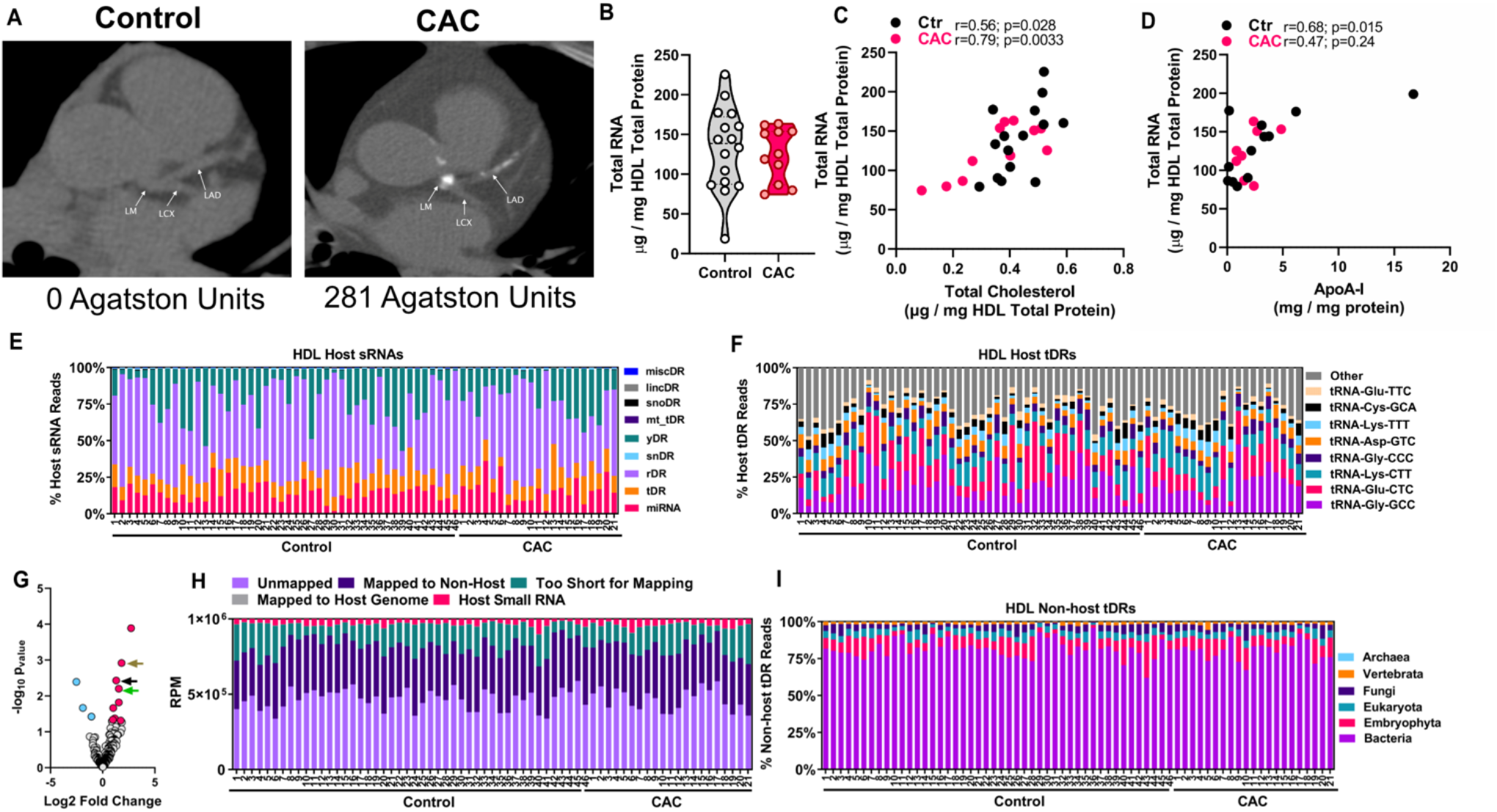
Distribution of small non-coding RNAs on human HDL. **A**) Non-contrast CT (computed tomography) scans from representative Control and individual with calcified coronary atheroma and CAC score >0. Left, CAC=0; right, CAC=281. Arrows indicate LAD (left anterior descending coronary artery), LCX (left circumflex artery) and LM (left main coronary artery). **B**) Total RNA content as determined by direct SYTO RNASelect staining (50 μM) on DGUC-HDL from Control (n=16) and CAC subjects (n=11), normalized to total protein. Violin plot showing median and quartile ranges. **C**) Correlation of Total RNA content from (B) and total cholesterol levels normalized to total protein. **D**) Correlation of Total RNA content (B) and apoA-I protein (ELISA) normalized to total protein. Results from sRNA-seq analysis of DGUC-HDL in Control (n=46) and CAC (n=21) subjects. **E**) Distribution of host sRNAs abundance as percentage of total host reads. **F**) Percentage of host tDR by anticodon. **G**) Differentially altered tDRs reads. Altered tDR-GlyGCC reads indicated with arrows, brown=tDR-GlyGCC-30; black=tDR-GlyGCC-33; green=tDR-GlyGCC-32. **E**) Reads per million (RPM) identified as host, non-host, genome, unmapped, or too short for conservative mapping with TIGER [8]. **F**) Alignment to non-host (nonhuman) tDR database grouped by taxa.

Host miRNAs are only a minor fraction of HDL-sRNA cargo and further investigation of non-miRNA classes for content and function are warranted. tDRs are associated with cellular stress and likely function as sRNA regulators of cell survival and protein synthesis[36–45]; however, the functional impact of extracellular HDL-tDRs remains to be determined. To define the human HDL-tDR profile and quantify differences in HDL-tDR levels in atherosclerosis (CAC vs Control), sRNA-seq and informatics (TIGER pipeline) were performed[8]. Pre-processed reads from HDL-sRNA-seq were aligned to the human reference genome (hg19) as well as to additional parent tRNA transcripts not present in the reference genome[8]. Special considerations were made for tRNA nomenclature, multi-mapping, and terminal 3’ CCA masking; and tDR read counts were tabulated for parent tRNAs at the amino acid anticodon level, as well as individually at the read level[8]. The most abundant class of host (human) sRNAs on DGUC-HDL were rDRs, followed by yDRs (Y RNA derived sRNAs), miRNAs, and tDRs (**Fig.1E**). Differential expression analyses (DEseq2) were used to identify altered HDL-sRNAs and multiple miRNAs, rDRs, and tDRs were found to be significantly increased or decreased on HDL from human atherosclerotic subjects (CAC) compared to control subjects (**Fig. S1C**). HDL acceptance of tDRs from stressed-donor cells may relieve stress-associated tDR-mediated gene regulation. Conversely, HDL-tDR delivery to recipient cells within the atherosclerotic lesion may contribute to cell-to-cell communication networks in atherosclerotic cardiovascular disease (ASCVD). Summary counts for parent tRNA-GlyGCC were the highest on human HDL, followed by tRNA-GluCTC and tRNA-LysCTT (**Fig. 1F**, **Table S3**). Multiple tDRs were observed to be significantly altered in human atherosclerosis at the individual sRNA level, as opposed to the parent transcript level (**Table S4**). For example, HDL levels of tDR-GlyGCC-30, a 30nt tDR processed from the 5’ half of parent tRNA-GlyGCC, were increased 3.5-fold (p=0.0012) on HDL from CAC subjects compared to control subjects (**Fig. 1G, Table S4**).

In our previous study, mouse HDL were demonstrated to transport many non-host sRNAs, including microbial sRNAs from bacteria and fungi[8]. To determine if human HDL also transport microbial sRNA, all non-host reads were further aligned to curated non-host genomes, including bacteria, fungi, and viruses. Strikingly, we found that DGUC-HDL from human plasma are also highly-enriched with non-host microbial sRNAs (**Fig.1H**). To determine if HDL transports non-human tDRs, non-host reads were further aligned to a database of tRNA transcripts (GtRNAdb), and human HDL were observed to carry many bacterial tDRs (**Fig.1I**). These results support the notion that human HDL transport high levels of sRNA in circulation, and HDL-sRNA levels are correlated to HDL-cholesterol and apoA-I levels. Moreover, results support the notion that human HDL transport host and non-host (bacterial) tDRs which are altered in atherosclerosis; however, the mechanism by which HDL binds to extracellular tDRs is unknown.

### Immune cell export of tDRs to HDL (Fig. 2)

We have previously observed that macrophages and other immune cells readily export miRNAs to HDL *in vitro*[9, 21]; however, it is unknown if macrophages also export other sRNA classes to HDL. Cellular tDR changes are associated with stress and macrophages within the atherosclerotic lesion are under both environmental (local tissue inflammation) and metabolic stresses (cholesterol-loading endoplasmic reticulum stress)[37, 46]. To determine if macrophages export tDRs to HDL, CD14+ human peripheral blood mononuclear cells (PBMC) were isolated from blood, differentiated to macrophages for 7d with GM-CSF (50ng/mL), and incubated with 1 mg/mL DGUC-HDL for 24h in serum-free media (**Fig. 2A**). HDL were repurified from culture media by SEC and total RNA was isolated from equal protein levels (0.2mg HDL total protein) of HDL pre- and post-incubation with macrophages. To quantify tDR-GlyGCC-30 export, qPCR with custom probes were performed and we observed an increase in HDL-tDR-GlyGCC-30 levels on post-HDL compared to pre-HDL (**Fig. 2B**). To confirm macrophage HDL-tDR export, mouse bone marrow cells were differentiated to bone marrow derived macrophages (BMDM) with GM-CSF for 7d, transiently-transfected for 24h with fluorescence-labelled tDR-GlyGCC-30-AF647, washed, and incubated with accepting HDL (1mg/mL) for 24h (**Fig. 2C**). BMDM culture media was collected and separated by SEC, and HDL-tDR-GlyGCC-30 export was determined by the levels of fluorescence (tDR-GlyGCC) in HDL (SEC) fractions. We observed a significant increase in fluorescence signal in fractions corresponding to HDL compared to media from wells that did not receive accepting HDL (**Figs. 2D, E**). These results suggest that primary macrophages in both humans and mice have the capacity to export tDRs, namely tDR-GlyGCC-30, to HDL *in vitro*. Although the mechanism by which HDL accepts tDRs from macrophages is unknown, HDL may have the capacity to bind to free, unprotected extracellular tDRs exported from macrophages.

**Figure 2.**
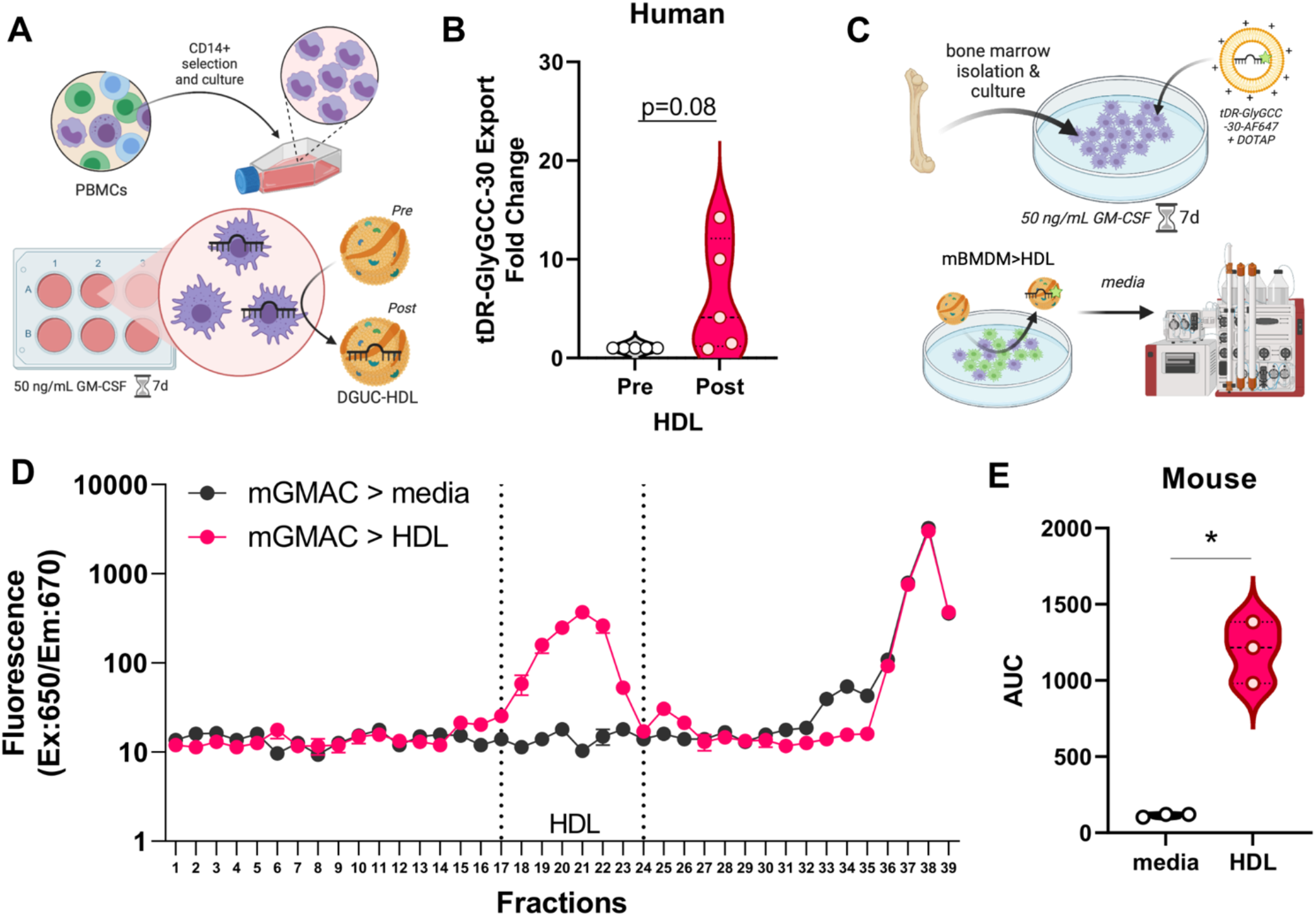
Macrophage export of sRNAs to HDL. **A**) Schematic representation of experimental design used in (B). **B**) HDL tDR-GlyGCC-30 expression by real-time PCR before (Pre) and after (Post) exposure to human GM-CSF differentiated macrophages. Violin plot showing median and quartile ranges, n=5. **C**) Schematic representation of experimental design used in (D-E). **D**) Fluorescence signal of exported tDR-GlyGCC-30 (AF647) from mouse bone marrow derived macrophages (mGMAC) to serum free media (mGMAC>media) or DGUC-HDL (mGMAC>HDL) across SEC fractions (1.5mL), n=3. **E**) Area under the curve (AUC) of HDL region from (D). Violin plot showing median and quartile ranges. *p<0.05. Schematics created with BioRender.com.

### Differential sRNA binding affinities to HDL (Fig.3)

To quantify the binding affinity of sRNAs to HDL, microscale thermophoresis (MST) assays were completed with oligonucleotides labeled with a 3’terminal fluorophore (Alexa Fluor 647) (**Figs. 3A, B**, **Table S2**). As a positive control, the binding affinity of tDR-GlyGCC-30 to its complementary antisense (AS) sequence was tested and strong affinity towards AS-GlyGCC-30 was observed (**Fig. 3C**). tDR-GlyGCC-30 binding to a control scrambled sequence (AS-Scr) was also assayed as a negative control, and tDR-GlyGCC-30 binding was not observed (**Fig.3C**). To calculate HDL’s binding affinity to tDR-GlyGCC-30, multiple DGUC-HDL samples were tested with MST and we found that HDL does indeed have strong affinity towards tDR-GlyGCC-30 (**Fig.3D**). To determine if HDL has similar affinity to other tDRs of same length (30nts), MST assays were used to calculate HDL affinities towards tDR-LysCTT-30 and tDR-GlyCCC-30, both of which were found to have reduced affinity towards HDL compared to tDR-GlyGCC-30 (**Fig.3E**). These results suggest that HDL have preferential binding towards tDR-GlyGCC-30 compared to other HDL-tDRs. Based on informatics, HDL-sRNAs are likely singlestranded RNA (ssRNA); however, tDR-GlyGCC has been reported to form secondary hairpin structures that have been proposed to increase its stability and resistance to RNases in plasma[47, 48], but could also influence HDL association. To determine if HDL preferentially bind to ssRNA or double-stranded RNA (dsRNA), dsRNA-tDR-GlyGCC-30 was synthesized in ssRNA and dsRNA forms (**Table S2**) and tested with MST. Human DGUC-HDL again showed strong affinity to ssRNA-tDR-GlyGCC-30; however, HDL failed to bind to dsRNA-tDR-GlyGCC-30, suggesting that HDL may indeed prefer ssRNA over dsRNA, at least for tDR-GlyGCC-30 (**Fig.3F**). Denaturing dsRNA-tDR-GlyGCC-30 prior to MST with heat to break apart the duplex was found to restore HDL binding (**Fig.3F**). Parent tRNAs can also be cleaved to produce multiple forms of sRNAs originating from the 5’ or 3’ terminal ends, often referred to as tRNA-derived halves (tRH) or tRNA-derived fragments (tRF)[37, 38, 49, 50]. Our candidate, tDR-GlyGCC-30, is a 5’-tRH likely cleaved by angiogenin in response to stress[37, 42, 44]; however, other forms and lengths of tDR-GlyGCC can also be cleaved. To determine if HDL have strong affinity to other potential tDRs arising from the parent tRNA-GlyGCC, specific tDR-GlyGCC fragments were synthesized and tested for binding affinity using MST (**Table S2**). HDL bound strongly to all candidate tDR-GlyGCC of variable lengths and sequences; however, the strongest binding affinity for HDL was observed to be our candidate tDR-GlyGCC-5’tRH-30, compared to tDR-GlyGCC-3’-tRF-15, tDR-GlyGCC-3’tRH, and tDR-GlyGCC-5’tRF-15 (**Figs.3G, H**). These results agree with observations from sRNA-seq of human HDL, as positional alignment analyses of tDRs mapped to their parent tRNAs showed enrichment for 5’ tRHs over 3’ tRHs for GlyGCC (**Fig.S2A**).

**Figure 3.**
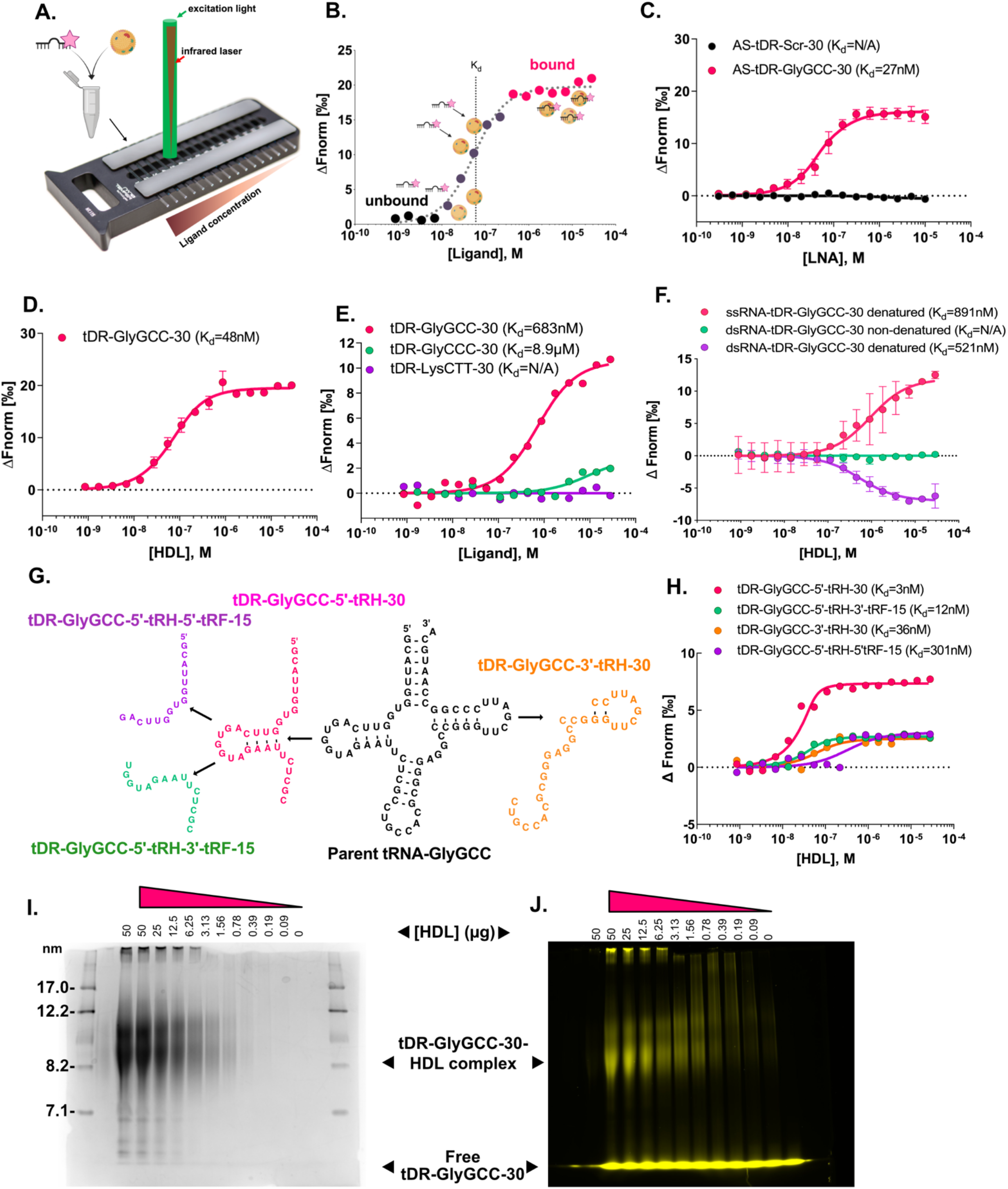
tDR-GlyGCC binding affinity to HDL by microscale thermophoresis. **A**) Setup of microscale thermophoresis (MST). Titrated ligand (e.g. DGUC-HDL) and fluorescently labelled (AF647) target (e.g. single stranded sRNA oligos) were combined (1:1) in glass capillaries and loaded onto the apparatus where an infrared laser heats the complex to create a small temperature gradient. sRNAs with HDL will demonstrate slower movement compared to free sRNA molecules. The movement of the fluorescent sRNA is then tracked. **B**) Example of a binding curve depicting difference between unbound sRNAs and bound sRNA molecules thermophoretic movement, where Fnorm is the normalized fluorescence against the ligand concentration, ΔFnorm is the baseline corrected normalized fluorescence. Data were fitted to the MST (K_d_) model. **C**) MST analysis of interactions between tDR-GlyGCC-30 and a scrambled antisense (AS) oligonucleotide (AS-Scr) or a complementary antisense to tDR-GlyGCC-30 (AS-tDR-GlyGCC-30), n=3. **D**) MST binding curves of tDR-GlyGCC-30 and DGUC-HDL, n=6. **E**) Representative MST binding curves of tDR-GluCTC and **E**) miR-223-3p compared to tDR-GlyGCC. Representative MST analysis of interactions between DGUC-HDL and tDR-GlyGCC-30, tDR-GlyCCC-30, or tDR-LysCTT-30. **F**) Binding curves of DGUC-HDL to single stranded (ssRNA) or double stranded (dsRNA) tDR-GlyGCC-30 without (non-denatured) or with heat (90°C 2min) denaturing (denatured), n=2-3. **G**) Schematic of tDR-GlyGCC fragments synthesized used in (H). **H**) Representative MST analysis of interactions between DGUC HDL and tDR-GlyGCC fragments in (G). Binding affinities (K_d_) are shown. Electrophoretic mobility shift assay (EMSA) with fluorescently (AF546) labelled sRNA and DGUC-HDL. tDR-GlyGCC (AF546) was incubated with titrations of HDL (1:1) and crosslinked (@254nm) for 15 min. Samples were loaded onto a 4-16% NativePAGE gel with 0.5X TBE running buffer. **I**) Coomassie of non-denaturing gel. Stokes radius (R_s_) of proteins standards shown. **J**) Fluorescent (AF546) migration of tDR-GlyGCC-30 on gel with free oligo and oligo complexed to HDL titrations is shown. Schematics created with BioRender.com.

In addition to MST, EMSA were used as an orthogonal method to demonstrate HDL’s binding affinity to tDR-GlyGCC-30. Briefly, tDR-GlyGCC-30-AF546 was cross-linked (UV, 254nm) to serial dilutions of DGUC-HDL for 15min. Bound and unbound tDR-GlyGCC-30/HDL complexes were separated by native polyacrylamide gel electrophoresis, and RNA binding was visualized with a fluorescence scanner (**Figs. 3I, J**). No fluorescence/RNA signal was observed in the HDL only lane; however, tDR-GlyGCC-30-AF546 crosslinked to HDL resulted in a shift (increase) in an HDL size (bands between: 8.2-12.2nm, **Fig. 3I, J**) in a dose-dependent manner. Moreover, the band position of HDL-tDRGlyGCC-30 complex was not observed in the RNA only lane. Based on HDL titrations, the binding affinity of HDL to tDR-GlyGCC-30-AF546 was calculated (**Fig. S2B**), as previously described [32], and results from EMSA confirmed that HDL harbors strong affinity towards tDR-GlyGCC-30. These results support that HDL preferentially bind to ssRNAs and possess strong affinity to tDRs based on length and sequence.

### Biochemical properties of sRNAs that impact HDL binding (Fig.4)

To calculate the binding affinity of DGUC-HDL to other classes of sRNAs present on HDL, candidate miRNAs were tested with MST. We have previously reported that miR-223-3p and miR-375-3p are abundant on circulating HDL[9, 21, 22]. MST assays here confirmed that HDL binds to both miR-223-3p and miR-375-3p with strong affinities (**Figs. 4A, B**). To further understand the biochemical and physical properties that influence HDL’s association to oligonucleotides, we examined the 2’ position of the ribose in ribonucleotides as this position distinguishes RNA from DNA. To determine the affinity of HDL to DNA, which lacks a hydroxyl group at the 2’ position, single-stranded DNA (ssDNA)-tDR-GlyGCC-30 was synthesized (**Table S2, Fig.4E**) and tested for HDL binding with MST. Results showed that HDL readily bound to the ssRNA form but failed to bind to the DNA form of tDR-GlyGCC-30 (**Fig.4C, D, E**). ssDNA-tDR-GlyGCC-30 harbored DNA bases along the length of the tDR-GlyGCC oligonucleotide, including thymines in lieu of uracils at 10 positions (**Table S2, Fig. 4E**). These results suggest that HDL binding to sRNA is enhanced by either the 2’ hydroxy group present on RNA or by uracil present in sRNA sequences. To determine if uracil facilitates sRNA binding to HDL, ssRNA form of tDR-GlyGCC-30 was synthesized with rU>T mutations at regional positions along the oligonucleotide. Mutations were created at the 5’ terminal end (5’T), 5’ terminal end + middle (5’midT), or the entire oligonucleotide (5’mid3’T) (**Table S2, Fig. 4E**). Based on MST assays, each of these three designs were observed to bind to HDL; however, the 5’T harboring the least amount of thymines showed the strongest binding affinity towards HDL. HDL’s affinity towards the 5’T mutant form of tDR-GlyGCC-30 was less than the native form of tDR-GlyGCC-30 (**Fig.4F**). These results support that sRNA uracil content is associated with increased binding affinity to HDL.

**Figure 4.**
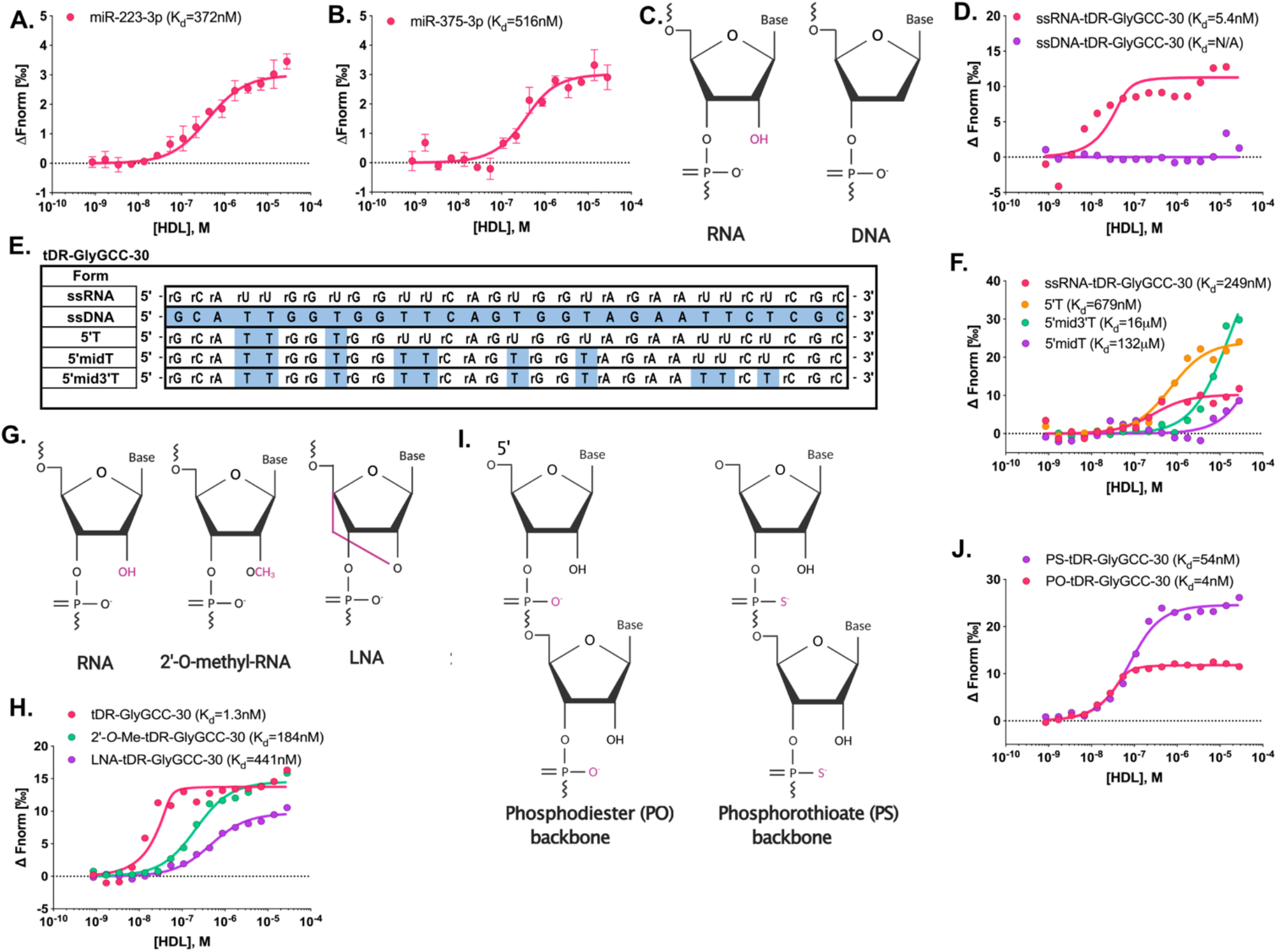
Biochemical properties of tDR-GlyGCC binding to HDL using MST. MST analysis of interactions between DGUC-HDL and **A**) miR-223-3p, n=3, or **B**) miR-375-3p, n=3. **C**) Structure of ribonucleotides (RNA) and deoxyribonucleotides (DNA). Hydroxyl group on RNA highlighted. **D**) Representative MST binding curves of single-stranded RNA (ssRNA-) or singlestranded DNA (ssDNA-) of tDR-GlyGCC-30. **E**) Synthesized tDR-GlyGCC-30 sequences of ssRNA, ssDNA, and rU>T mutations: at the 5’ terminal end (5’T), the 5’ terminal end and middle (5’midT), and the entire oligonucleotide (5’mid3’T). **F**) Representative MST binding curves of DGUC-HDL with tDR-GlyGCC-30 and its rU>T mutations from (E). **G**) Structure of RNA with 2’-O-methylation (2’-O-Me) or locked-nucleic acid (LNA) which harbors a bridge between 2’ and 5’ positions of the base ring. Modifications highlighted. **H**) Representative MST binding curves of DGUC-HDL and tDR-GlyGCC-30 with either 2’-O-Me or LNA modifications. **I**) Structure of RNA with phosphodiester (PO) or phosphorothioate (PS) backbones. **J**) Representative MST binding curves of DGUC-HDL and tDR-GlyGCC-30 with either PO or PS backbones. Binding affinities (K_d_) are shown. Structures created with BioRender.com.

To investigate the impact of the 2’ position on HDL-sRNA binding, tDR-GlyGCC-30 was synthesized with an O-linked methyl group at the 2’ position in place of the normal hydroxyl group, and MST assays found that 2-O-methylation reduced HDL binding affinity (**Fig.4G, H, Table S2**). Locked-nucleic acids (LNA) contain a bridge between the 2’ and the 5’ positions of the base ring, which locks the oligonucleotide in rigid confirmation and disrupts the 2’ ribose position (**Fig.4G**). To determine if tDR-GlyGCC-30 harboring LNA bases (instead of a normal hydroxyl group) reduces HDL-sRNA affinity, MST assays were completed. Based on MST, HDL were found to have decreased affinity towards LNA-tDR-GlyGCC-30 compared to the native form of tDR-GlyGCC-30 (**Fig.4H**). These results support that the 2’ hydroxyl promotes sRNA binding to HDL as disruption of this position through two different approaches both reduced HDL-sRNA binding affinity. Natural sRNAs have phosphodiester (PO) bonds (**Fig.4I**) between nucleosides which are sensitive to RNase hydrolysis. To increase sRNA stability for experimental studies, oligoribonucleotides are often synthesized with phosphorothioate (PS) bonds (**Fig.4I**) to increase resistance to RNases digestion, e.g. siRNAs for gene silencing [51]. Based on MST analyses, HDL were found to bind to both PO and PS forms of tDR-GlyGCC-30 with strong affinities (**Fig.4J**). Collectively, these results demonstrate critical features of sRNA biology that influence HDL association.

### Apolipoprotein A-I is a sRNA binding protein (Fig.5)

To determine if tDR-GlyGCC-30 has similar binding affinities to other lipoproteins, apolipoprotein B-containing lipoproteins (VLDL and LDL) were isolated by DGUC and tested with MST. Due to the molecular mass of the VLDL, binding affinities were calculated as EC50, and both VLDL and LDL were found to bind to tDR-GlyGCC-30 with greater affinity than HDL (**Fig.5A**). Based on tDR-GlyGCC-30’s strong affinity to all three lipoprotein classes, we sought to determine if sRNAs bind to the phospholipid or protein components of HDL. We first assessed if the RNA cargo separates with lipid or protein cargo by delipidation studies. HDL were isolated from human plasma by DGUC and delipidated using the Bligh-Dyer method to separate phospholipids away from protein cargo. Total RNA levels were quantified in input DGUC-HDL and matched delipidated total protein post-Bligh-Dyer lipid extraction by SYTO RNASelect, and the levels of detectable total RNA on remaining HDL samples were not reduced by phospholipid removal (**Fig.5B**). In addition, apoA-I was further isolated from the delipidated total protein samples by ion exchange separation (on Akta Pure FPLC system) and tested for sRNA content, which showed that apoA-I retained high levels of total RNA after delipidation of the HDL particles and ion exchange separation of apoA-I from other HDL proteins (**Fig.5B**). These results suggest that sRNAs likely associate with the protein component of HDL, namely apoA-I, as opposed to HDL phospholipids. To determine if apoA-I is a novel RNA binding protein, human apoA-I was purified from DGUC delipidated HDL using ion exchange columns and tested for binding affinity to tDR-GlyGCC-30 with MST. ApoA-I was found to bind to tDR-GlyGCC-30 with moderate affinity, confirming that apoA-I is likely an RNA binding protein (**Fig.5C**). As a negative control, actin (isolated from rabbit muscle) was observed to have very weak binding to tDR-GlyGCC-30 by MST (**Fig.S3A**). In addition to tDR-GlyGCC-30, apoA-I was also found to bind to tDR-GluCTC-30, miR-223-3p and miR-375-3p (**Fig.S3B-D, Table S2**). To determine if tDR-GlyGCC binds to phospholipids, PC-based single unilamellar vesicles (SUV) were generated and tested with MST. All sRNAs tested completely failed to bind to phospholipid SUVs, including tDR-GlyGCC-30, tDR-GluCTC-30, miR-223-3p and miR-375-3p (**Figs.5D, S3E-G**). These findings indicate that extracellular sRNAs do not directly bind to phospholipids on lipoproteins. Since sRNAs were observed to bind to native HDL (**Fig.3**) with greater affinity than lipid-free apoA-I (**Fig.5B**), MST assays were used to quantify sRNA binding to reconstituted HDL (rHDL), and tDR-GlyGCC-30 was observed to have moderate binding affinity to rHDL with apoA-I (rHDL+apoA-I); however, tDR-GlyGCC failed to bind to rHDL-like lipid particles when apoA-I was not present on rHDL (rHDL-apoA-I) (**Fig.5E**). To determine if sRNAs bind to generic lipoproteins or protein—lipid complexes mimicking HDL, synthetic HDL (sHDL) were generated through reconstitution of synthetic apoA-I mimetic peptides with distinct phospholipids shells. For example, ESP-24218, a synthetic apoA-I mimetic peptide consisting of 22 amino acids[52], was reconstituted with 1-palmitoyl-2-oleoyl-sn-glycero-3-phosphocholine (POPC) to form ESP-24218-POPC-sHDL. Strikingly, ESP-24218—POPC-sHDL failed to bind to tDR-GlyGCC-30 in MST assays (**Fig.5F**). To test if tDR binds to ESP-24218-sHDL composed of other phospholipid species, sHDL were reconstituted individually using 1,2-dimyristoyl-sn-glycero-3-phosphocholine (DMPC), 1,2-dipalmitoyl-sn-glycero-3-phosphocholine (DPPC), 1,2-distearoyl-sn-glycero-3-phosphocholine (DSPC), 1,2-dipalmitoyl-sn-glycero-3-phosphorylglycerol (DPPG), or 1,2-Dimyristoyl-sn-glycero-3-phosphoethanol-amine (DMPE). Despite altering the phospholipid chain lengths and head groups, sHDL particles were not observed to bind to sRNAs by MST analyses (**Fig.S4A-E**). These results support that apoA-I is a novel RNA binding protein for sRNAs, including tDRs and miRNAs. Moreover, results indicate that apoA-I and not phospholipids confer HDL particle binding and association to extracellular sRNAs.

**Figure 5.**
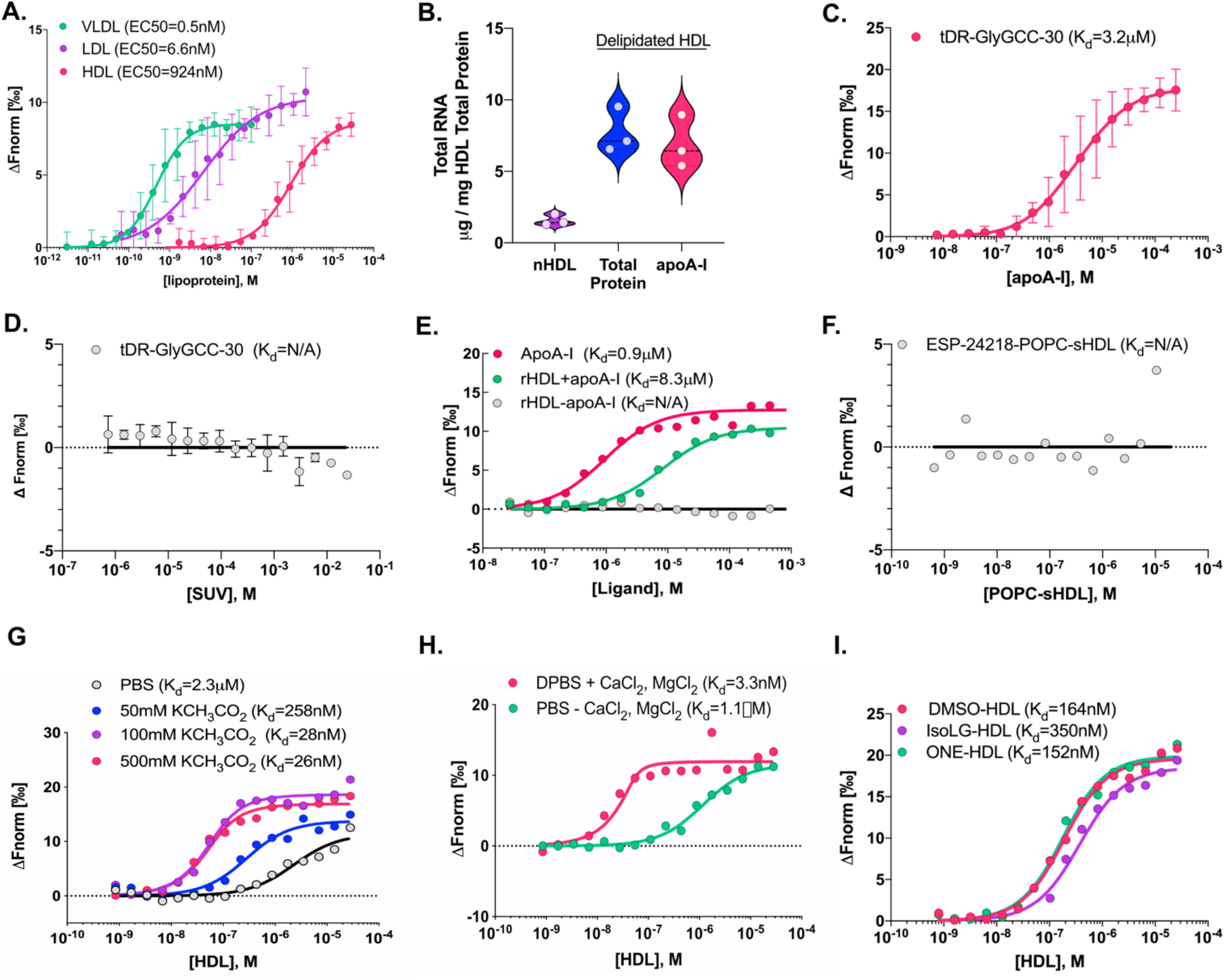
tDR-GlyGCC binds to apolipoprotein A-I. **A**) MST analysis of interactions with tDR-GlyGCC-30 and DGUC-HDL, -LDL, or -VLDL, n=3. **B**) Total RNA content as determined by direct SYTO RNASelect staining (50 μM) on matched human DGUC-HDL (nHDL), delipidated DGUC-HDL (total protein) and (FPLC) purified apoA-I from delipidated DGUC-HDL. Violin plot showing median and quartile ranges, n=3. **C**) MST analysis of interactions with tDR-GlyGCC-30 and human DGUC-HDL purified apoA-I, n=3. **D**) MST analysis of interactions with tDR-GlyGCC-30 and small unilamellar vesicles (SUV), n=3. **E**) Representative MST binding curves of tDR-GlyGCC-30 binding to purified apoA-I, or reconstituted HDL (rHDL) with or without apoA-I (rHDL+/-apoA-I). **F**) Representative MST binding curve of tDR-GlyGCC-30 binding to synthetic apoA-I mimetic peptide (ESP-24218) reconstituted with 1-palmitoyl-2-oleoyl-sn-glycero-3-phosphocholine (POPC) to form ESP-24218-POPC-sHDL. **G**) Representative MST binding curves of tDR-GlyGCC-30 binding to DGUC-HDL with increasing concentrations of potassium acetate (KCH3CO2, 50-500mM) in the PBS buffer. **H**) Representative MST binding curves of tDR-GlyGCC-30 binding to DGUC-HDL with PBS or DPBS with calcium chloride (CaCl2) and magnesium chloride (MgCl2) in the buffer. **I**) Representative MST binding curves of tDR-GlyGCC-30 and DGUC-HDL treated with vehicle (DMSO), isoLG (isolevuglandins), or ONE (lipid aldehyde 4-oxo-2-nonenal). Binding affinities (K_d_) are shown.

To further investigate the biochemical properties of HDL-sRNA binding, multiple physico-chemical approaches were taken with MST assays. To determine if sRNA binding to HDL is ionic, MST assays were completed with human DGUC-HDL and tDR-GlyGCC-30 in a graded series of increasing potassium acetate concentrations. Remarkably, we found that HDL-sRNA binding affinities were increased with increasing concentrations of potassium acetate (**Fig.5G**). To further test the impact of salt conditions, HDL-sRNA binding was measured in the presence of 1X PBS and 1X DPBS (with CaCl2 and MgCl2, Corning) by MST, and results showed that HDL binding to tDR-GlyGCC-30 was greatly enhanced in the 1X DPBS (**Fig.5H**). These results indicate that HDL binding to sRNAs, e.g. tDR-GlyGCC-30, is not likely to be ionic, as the high salt concentrations increased binding affinity, and could suggest that the association between HDL and extracellular sRNA is hydrophobic. To further define chemical influences or regulators of HDL-sRNA binding, we sought to determine the impact of reactive modifications on the binding affinity between HDL particles and extracellular sRNAs, including isolevuglandin (IsoLG) and 4-oxo-nonenal (ONE) that modify positively charged residues of apoA-I (i.e. lysines and arginines) that have the potential to form ionic bonds with the negatively charged backbone of sRNAs. In ASCVD, HDL lose many of its beneficial functions, including cholesterol efflux acceptance capacity and reactive modifications to HDL’s lipids and protein (amino acids) likely confer these functional changes[53, 54]. We have previously reported that IsoLG-modifications to HDL particles are increased in ASCVD and associated with decreased cholesterol efflux capacity[55]. We have also found that ONE-modifications also reduce HDL cholesterol acceptance capacity[33]. To determine if IsoLG- or ONE-modifications also alter HDL’s ability to bind to sRNAs, DGUC-HDL was treated with 1 molar equivalents of IsoLG, ONE, or DMSO (vehicle control), and modified HDL were tested for binding with MST. Although these modifications alter HDL’s ability to accept cholesterol and other functions, they did not alter HDL’s ability to bind to tDR-GlyGCC-30 (**Fig.5I**), further supporting the notion that the association between HDL and extracellular sRNA is hydrophobic.

## DISCUSSION

HDL transport many different types of molecules, however, the mechanisms by which HDL interacts with non-lipid cargo are unknown. Here, we demonstrate that HDL binds to extracellular sRNAs through apoA-I, for which RNA binding is a previously unrecognized biological function. Results indicate that HDL carry high concentrations of both host and non-host tDRs, including tDR-GlyGCC-30. This tDR was highly abundant on HDL and was found to be significantly increased in subjects with advanced coronary atheroma as determined by the presence of raised calcified lesions by non-contrast cardiac CTs (CAC>0). Macrophages may contribute to the HDL-tDR signature as HDL were found to readily accept tDRs from primary macrophages. HDL-sRNA binding assays were completed by MST and confirmed by EMSA. These assays showed that HDL have strong affinity towards extracellular sRNAs, including tDRs and miRNAs, with dissociation constants (K_d_) in the nM range. To define the physico-chemical properties that regulate and influence HDL-sRNAs, tDR-GlyGG-30 was synthesized in multiple chemical forms, lengths, structures, and modifications to test in HDL binding assays. In this study, we found that the 2’ hydroxyl group present on nucleotide ribose is a strong determinant of HDL binding. For example, DNA (lacking the 2’-hydroxyl group), 2’-*O*-methylation of sRNA, and locked confirmation of the 2’ position, were each found to disrupt HDL-tDR binding. In subsequent experiments, we sought to determine the impact of specific reaction conditions on HDL-sRNA binding as a means to further understand the biochemistry of the association. We observed that high-salt conditions greatly increased HDL affinity towards tDR-GlyGCC-30, suggesting that the interaction between apoA-I and tDRs may not be solely through electrostatic interactions and may be conferred by hydrophobic interactions.

Results showed that human DGUC-HDL transport high concentrations of extracellular sRNAs in circulation and overall sRNA levels were correlated to total cholesterol levels in both control and atherosclerosis subjects (CAC). The use of SYTO RNASelect has allowed for the determination of total RNA content on HDL particles without the need to isolate RNA or have prior knowledge of candidate sRNAs for PCR quantification. Based on a ssRNA standard curve, we demonstrated that HDL transports approximately 125μg of RNA per mg of HDL total protein. This level of RNA is much greater than previously predicted or calculated based on quantitative PCR of individual miRNAs[27, 56, 57]. Based on sRNA-seq results, we also now recognize that host miRNAs are only a small percentage of the total amount of sRNAs on circulating HDL. The functional impact of HDL-sRNAs may not be restricted to a few select miRNAs, but most likely is conferred by the multitude of host and non-host sRNA classes, namely tDRs and rDRs, that make up the majority of sRNAs on HDL. The use of SYTO RNASelect dye here has illustrated the potential mass of regulatory sRNAs carried by HDL.

The most important advance of this study is the observation that HDL binds to sRNAs through apoA-I, not phospholipids. We have previously predicted that extracellular sRNAs may bind to HDL through binding to zwitterionic lipids (PC) within the phospholipid shell of HDL particles[9]. Oligonucleotides have been previously reported to bind to and associate with zwitterionic lipids, potentially through a divalent cation bridge between the positively charged choline headgroup on PC and the negatively charged backbone of RNA[24, 58, 59]. Others have expanded on this discussion towards a model in which an unknown RNA binding protein(s) on HDL, as opposed to phospholipids, confer the association of sRNAs to HDL particles[27]. In agreement with data from Plochberger and colleagues, our study clearly showed that phospholipid vesicles and/or particles did not bind to the sRNAs we tested whereas apoA-I protein did bind to multiple sRNAs. In this study, multiple approaches were used to determine if sRNAs bind to the protein or lipid component of HDL, and all evidence from this study supports that it is indeed a protein, not lipid, that mediates the association of sRNAs to HDL particles. For example, removing phospholipids from HDL particles did not change the amount of detectable RNA on HDL particles. Moreover, apoA-I, which accounts for approximately 70% of the protein mass on HDL, was demonstrated to be a novel RNA binding protein and bind to tDR-GlyGCC with moderate affinity, slightly less than HDL particles which showed strong affinity. tDR-GlyGCC-30 and other sRNAs were also found to bind to rHDL; however, construction of rHDL-like particles without apoA-I failed to produce particles that would bind to extracellular sRNAs. Collectively, these strongly suggest that sRNAs bind to the protein component, specifically apoA-I, and not the phospholipid component of HDL, as we have previously hypothesized[9].

Differential expression analysis identified many significantly altered tDRs on HDL in atherogenic subjects, including tDR-GlyGCC-30. The detection of tDR-GlyGCC in our HDL sequencing dataset is not uncommon, as this sequence has been detected in many biological samples, including both cellular and extracellular RNA samples[36, 37, 40, 47, 49, 60]. The underlying reason for tDR-GlyGCC high abundance in plasma may be related to increased stability and resistance to RNase digestion, as opposed to increased synthesis and secretion from donor cells[47, 48]. tDR-GlyGCC is likely detected in most extracellular sRNA-seq datasets due to its potential resistance to plasma RNases that degrade other secreted tDRs. Its protection from digestion may be related to its high potential to form homodimers with other tDR-GlyGCC or even heterodimers with tDR-GluCTC[47, 48]. Furthermore, tDR-GlyGCC may even form a small 6nt hairpin on its 5’ terminal end that may contribute to increased stability. For MST testing, all oligos were denatured with heat prior to analyses to prevent secondary structures; however, no differences were observed with tDR-GlyGCC-30 binding to HDL with or without heat denaturing prior to MST analysis (data not shown). We did observe, however, that HDL displayed preferential binding to ssRNA over dsRNA forms of tDR-GlyGCC, and HDL binding affinity to dsRNA-tDR-GlyGCC was restored when dsRNA-tDR-GlyGCC-30 was denatured (by heat) prior to MST analysis. These results support that while tDR-GlyGCC’s potential to form self-hairpins and/or homodimers may increase their stability in biofluids and increase the chance of associating lipoproteins, HDL’s affinity to tDR-GlyGCC is not likely affected by this potential secondary structure.

The biological functions of tDRs on HDL are unknown, but cellular tDRs have emerged as key regulators of cellular stress and protein synthesis[37, 41]. HDL have the capacity to deliver functional miRNAs to recipient cells, including endothelial cells[10, 12, 13]; however, HDL-tDR transfer to endothelial cells or other recipient cells has not been demonstrated to date. It’s plausible that HDL does transfer tDRs, like miRNAs, to recipient cells, and regulate gene expression, but these studies will require further investigation in the future. Currently, it is unknown if HDL can transfer sRNAs to immune cells, however, if HDL is taken up by immune cell phagocytosis or fluid-phase uptake, then HDL-tDRs cargo may activate ssRNA sensing Tolllike receptors 7/8 (TLR7/8) in immune cells, leading to pro-inflammatory gene expression, cytokine secretion, and inflammation. HDL is generally anti-inflammatory in health, but may promote inflammation in certain diseases or conditions[61]. HDL’s ability to bind and sequester extracellular tDRs, particularly non-host tDRs, likely serves as an anti-inflammatory function to prevent excess activation of pattern recognition receptors, i.e. TLR7/8, in immune cells. HDL also may regulate genes through acceptance of regulatory sRNAs away from donor cells. Here, we show that macrophages export tDRs to HDL; however, future studies are needed to define the impact of tDR export on macrophage gene expression. Likewise, future investigation is needed to determine if macrophages export tDRs to HDL in response to specific cellular stresses. Although macrophages export tDRs to HDL, the mechanism by which HDL acquires tDRs from macrophages is currently unknown. It is possible that macrophages and other cell types simply export unprotected and free sRNAs from the cell and HDL have the capacity to bind and protect free sRNAs. In this model HDL would interact with extracellular sRNAs independent of cells and would be entirely dependent on HDL’s affinity for sRNA. To calculate HDL’s binding affinity (K_d_, dissociation constant) to sRNAs and determine the underlying biochemistry of their association, MST and EMSA assay were used. Remarkably, HDL were found to have very strong binding affinity towards ssRNA forms of tDR-GlyGCC-30. The clinical applicability of these findings are unknown; however, therapeutic strategies based on HDL’s ability to bind to and delver RNAs to specific cells may ultimately elevate HDL as a potential RNA-based drug delivery vehicle. Conversely, results presented here may allude to a phenomenon in which HDL readily binds to extracellular sRNAs to sequester potentially inflammatory signals away from pattern recognition receptors. Future work is needed to define the role of HDL in dampening uncontrolled immune responses due to extracellular sRNAs; however, if established then HDL-based therapies may be applied to sepsis and infection.

Collectively, results support that HDL transport high concentrations of extracellular sRNAs, including both host and non-host tDRs. HDL was determined to bind to tDRs through the novel RNA binding protein apoA-I which also serves as the structure-function protein for HDL particles. HDL was found to bind to RNA over DNA and single-stranded over double-stranded forms. Key biochemical experiments showed that a hydroxyl group at the 2’ position of the ribose was determinative towards HDL binding and disruption of this group reduced HDL binding affinity to sRNAs. Likewise, HDL association to sRNAs might be mediated through hydrophobic interactions as HDL-sRNA binding was greatly increased in high salt buffers. Overall, results from our investigation of the underlying biochemistry of HDL’s association to sRNAs revealed key fundamental principles of HDL’s transport of tDRs in ASCVD and HDL’s affinity towards extracellular sRNAs. These results will serve as a platform for future studies to manipulate HDL-sRNA cargo for therapeutic approaches to prevent or treat ASCVD.

## DATA AVAILABILITY

All raw sequencing data will also be made available upon request.

## ACKNOWLEDGEMENTS

The authors would like to recognize Meaghan Kuzmich for assistance with lipoprotein procurement, Shilin Zhao for help with bioinformatics, Dr. Michelle Ormseth for assistance with statistical analyses, Dr. Loren Smith for clinical research methods and Angela Jones of VANTAGE at VUMC for expertise in high-throughput sequencing technologies.

## SUPPLEMENTARY DATA

**Figure S1.**
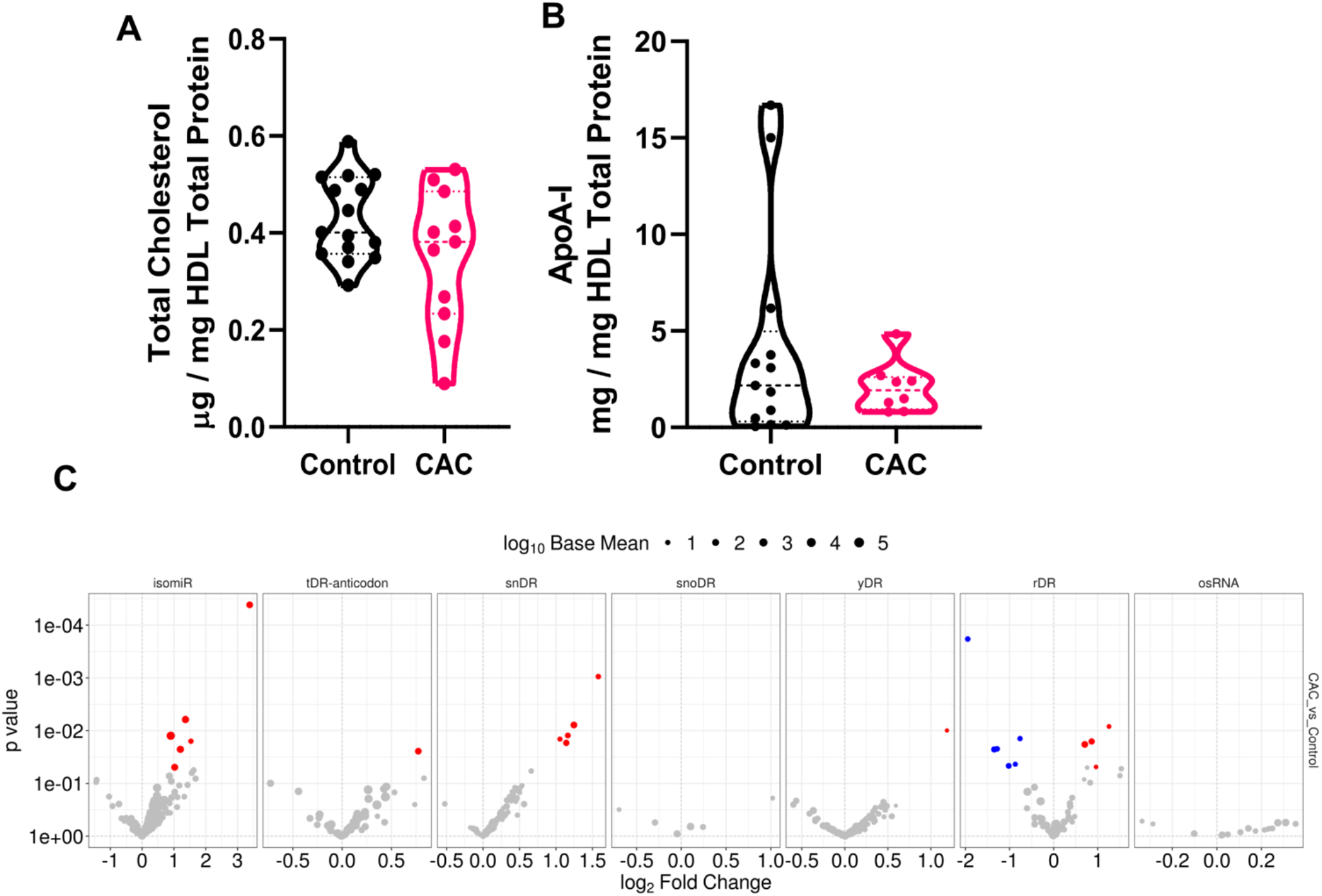
HDL isolated by density-gradient ultracentrifugation (DGUC) was assayed for **A**)total cholesterol and protein by colorimetric kit and **B**)ApoA-I protein by ELISA (n=11 Control, n=11 CAC sujects). Violin plots showing median and quartile ranges. **C**)Differential expression analysis at the parent level from sRNA sequencing of HDL from Control (n=46) and CAC (n=21) Subjects. Volcano plots demonstrating significant DESeq2 analysis and the log_2_ fold change. Red, increased; blue, decreased.

**Figure S2.**
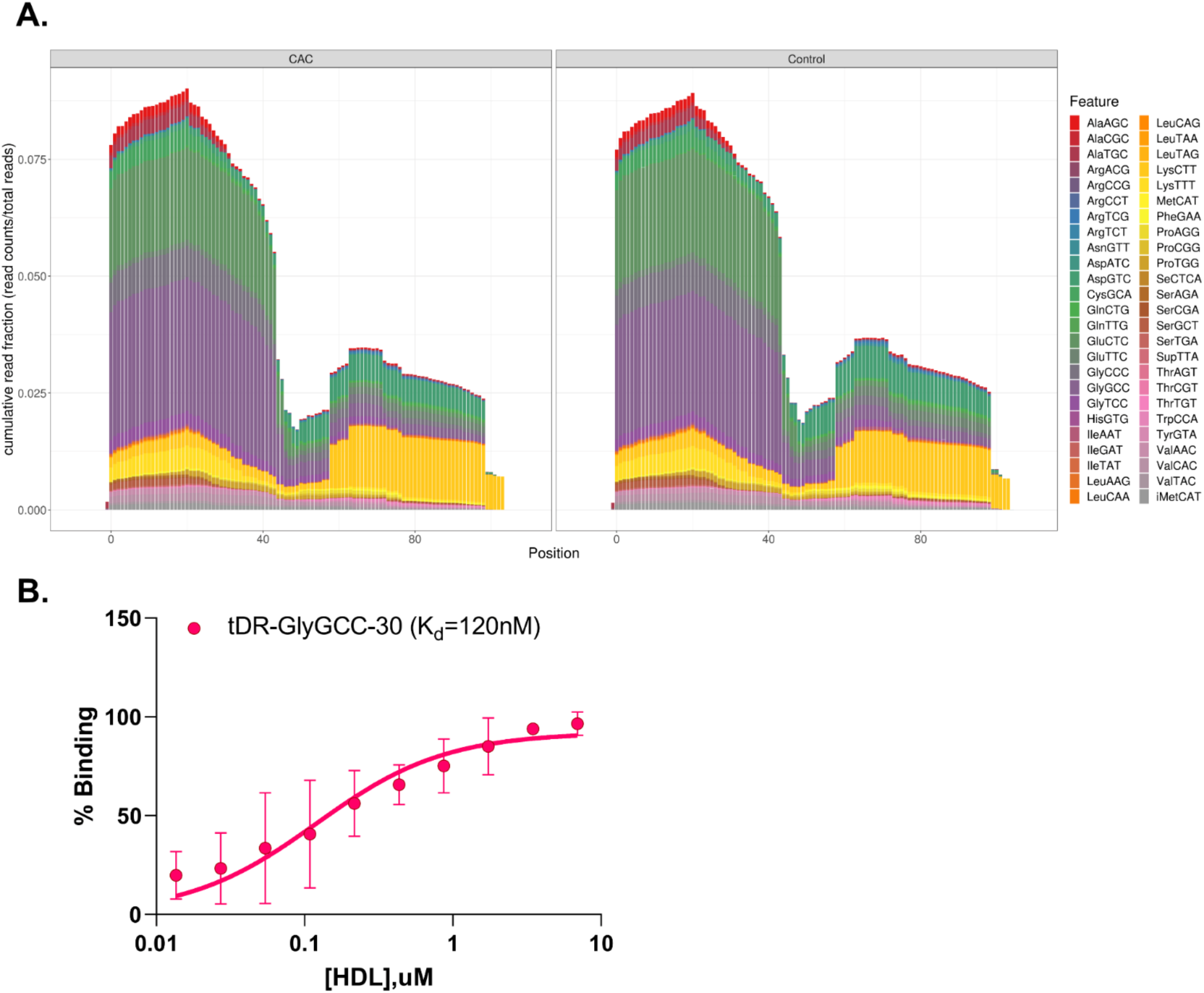
**A**) Positional coverage of maps of HDL tDRs for parent tRNA amino acid anticodons, reported as mean cumulative read fraction (read counts/total counts) between Control (n=46) and CAC (n=21) subjects. **B**) Fluorescence electrophoretic mobility shift assay nonlinear regression binding percentage of DGUC-HDL complexed with tDR-GlyGCC-30 labelled with Alexa Fluor 546. Binding affinity (K_d_) is shown. n=3, data are mean ± SD.

**Figure S3.**
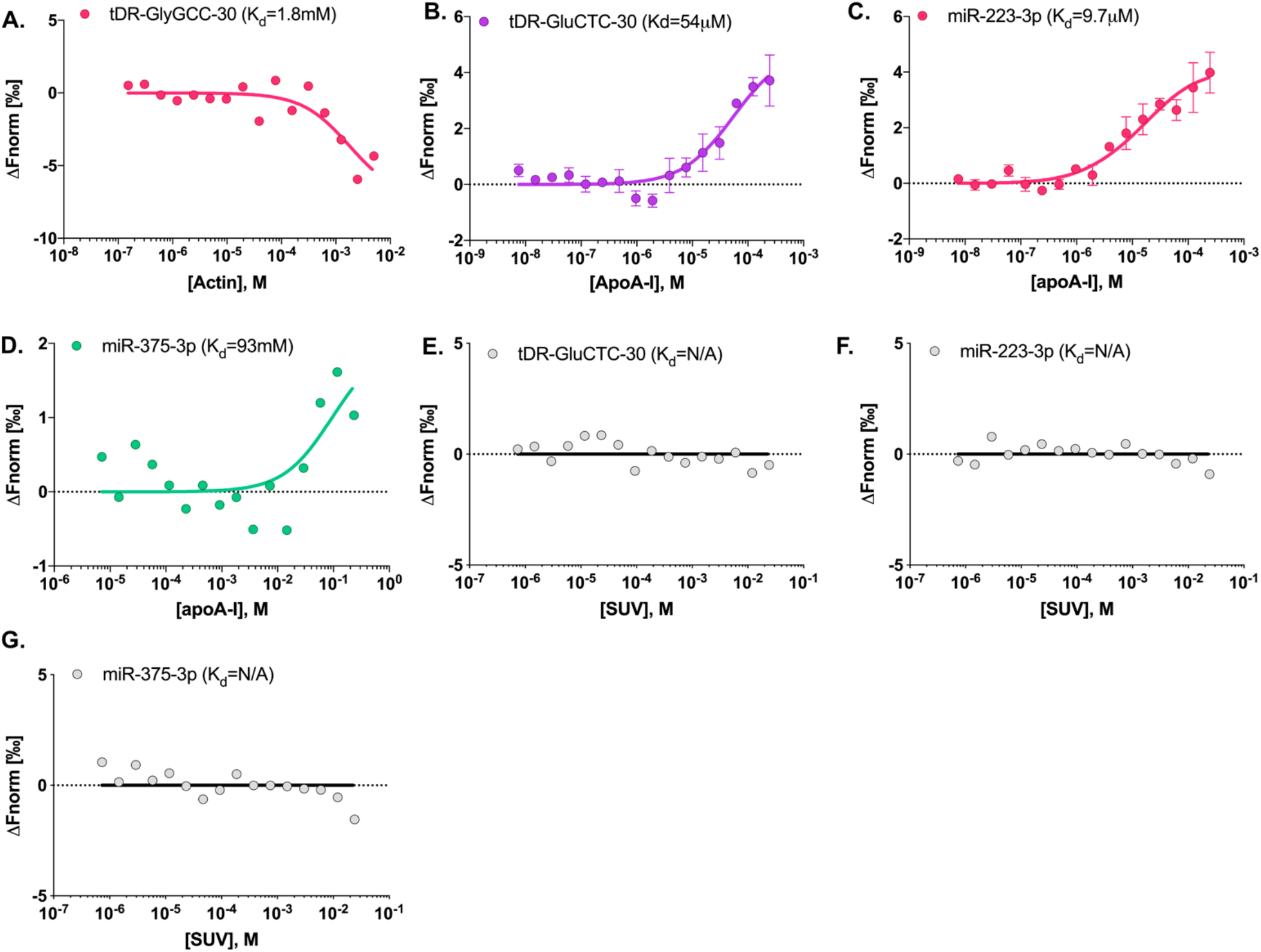
MST analysis of interactions between **A**) tDR-GlyGCC-30 and actin (rabbit muscle, Sigma), apoA-I and **B**) tDR-GluCTC-30, **C**) miR-223-3p, and **D**) miR-375-3p; and single unilamellar vesicles (SUV) and **E**) tDR-GluCTC-30, **F**) miR-223-3p, **G**) miR-375-3p. Binding affinities (K_d_) are shown.

**Figure S 4.**
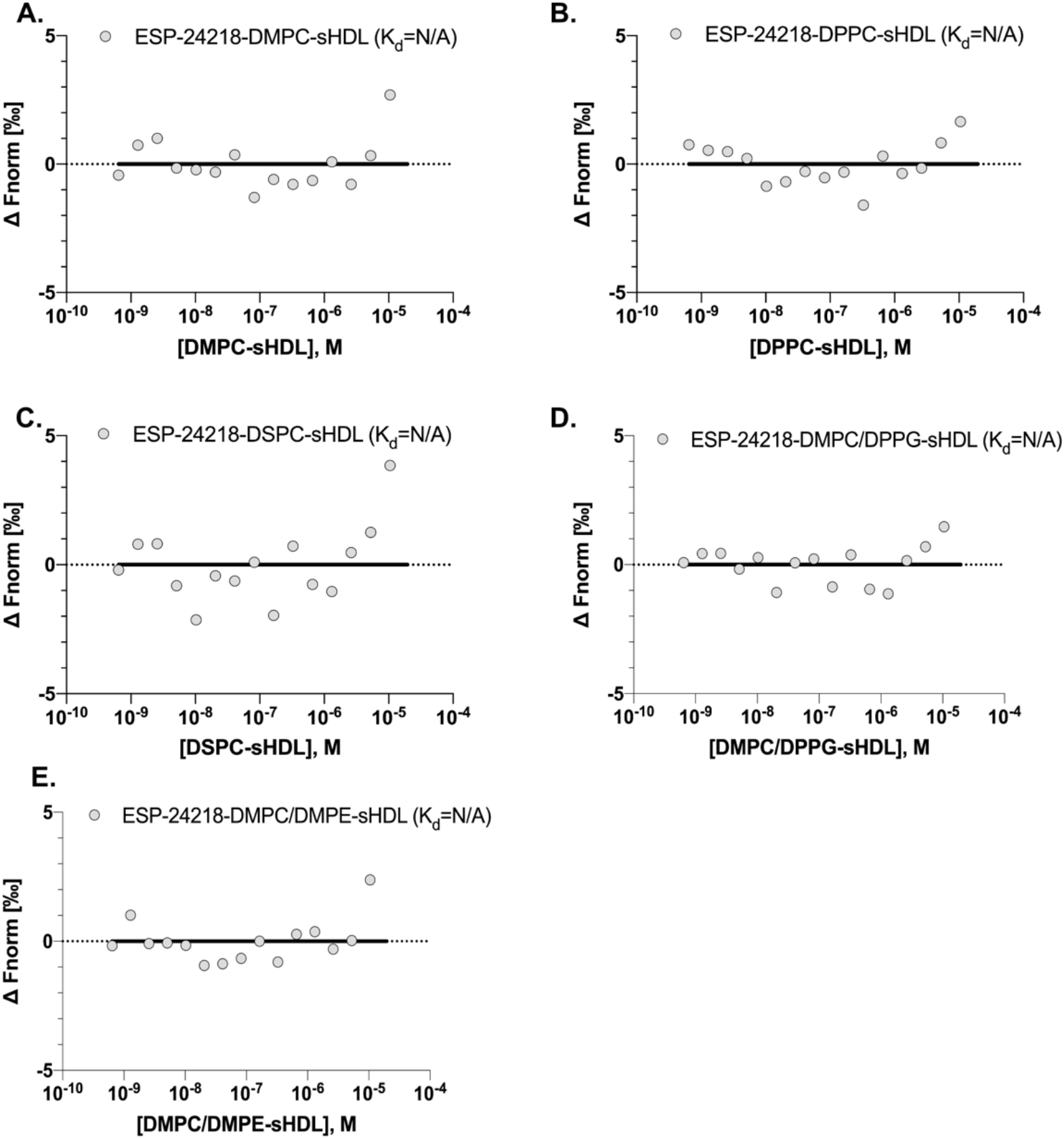
MST analysis of interactions between tDR-GlyGCC-30 and synthetic HDL (sHDL) with apoA-I mimetic peptide, ESP-24218, and either **A**) 1,2-dimyristoyl-sn-glycero-3-phosphocholine (DMPC), **B**) 1,2-dipalmitoyl-sn-glycero-3-phosphocholine (DPPC), **C**) 1,2-distearoyl-sn-glycero-3-phosphocholine (DSPC), **D**) DMPC and 1,2-dipalmitoyl-sn-glycero-3-phosphorylglycerol (DPPG), and **E**) DMPC and 1,2-Dimyristoyl-sn-glycero-3-phosphoethanol-amine (DMPE). Representative curves and binding affinities (K_d_) are shown.

**Table S1:** Clinical parameters for CAC and Control Subjects

|  | CAC (N=21) | Control (N=46) | P-value |
| --- | --- | --- | --- |
| Age, years | 61 (54, 67) | 45 (37, 56) | <0.001 |
| Race, # (% Caucasian) | 18 (86) | 39 (85) | 0.92 |
| Sex, # (% female) | 9 (43) | 30 (65) | 0.09 |
| Total-C, mg/dL | 218 (185, 326) | 191 (176, 267) | 0.14 |
| HDL-C, mg/dL | 53 (49, 74) | 57 (51, 67) | 0.91 |
| LDL-C, mg/dL | 142 (110, 221) | 118 (96, 182) | 0.23 |
| Triglycerides, mg/dL | 103 (73, 171) | 90 (76, 117) | 0.35 |
| Coronary calcium score, Agaston units | 61 (9, 221) | 0 (0, 0) | <0.001 |

Race, Sex, P value Pearson Chi-Square; P value Mann-Whitney U; Median (25% and 75% quartiles)

**Table S2.**
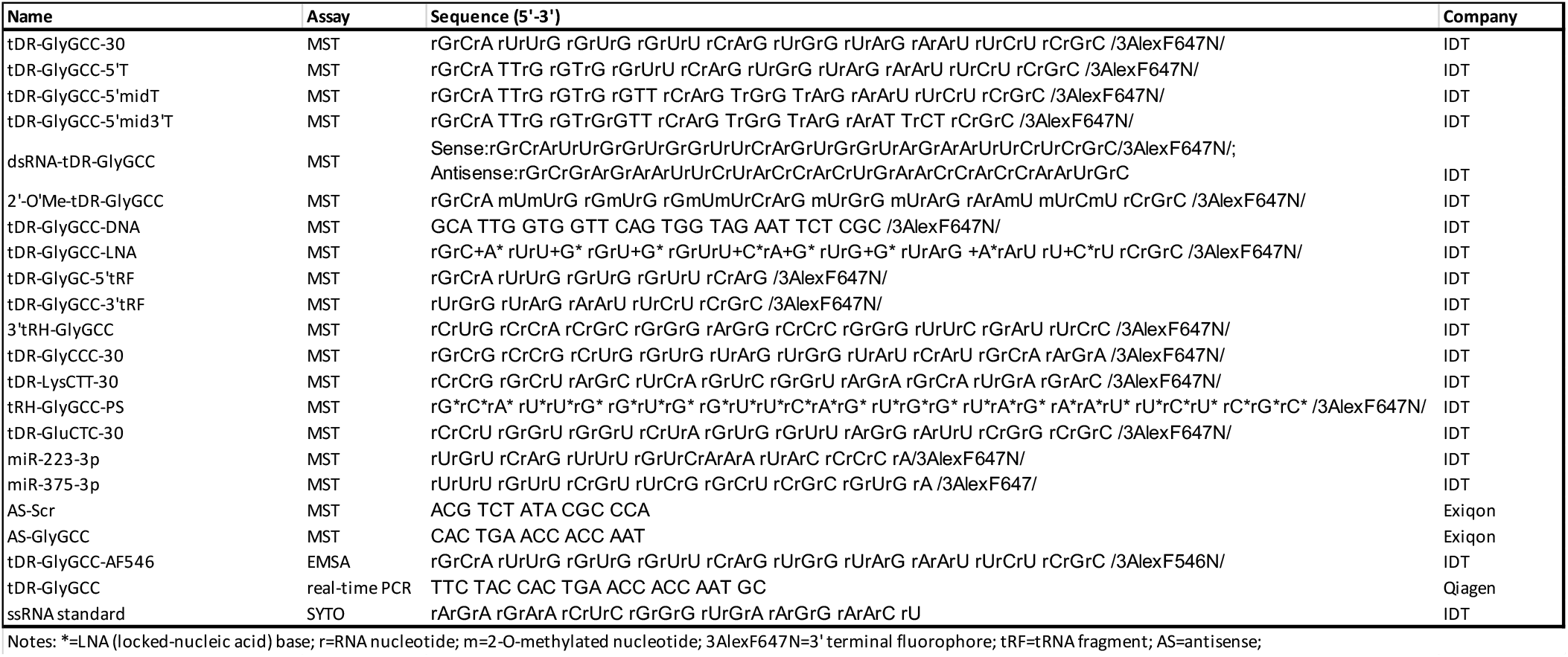
Sequences of synthesized oligonucleotides used in the study.

**Table S3.**
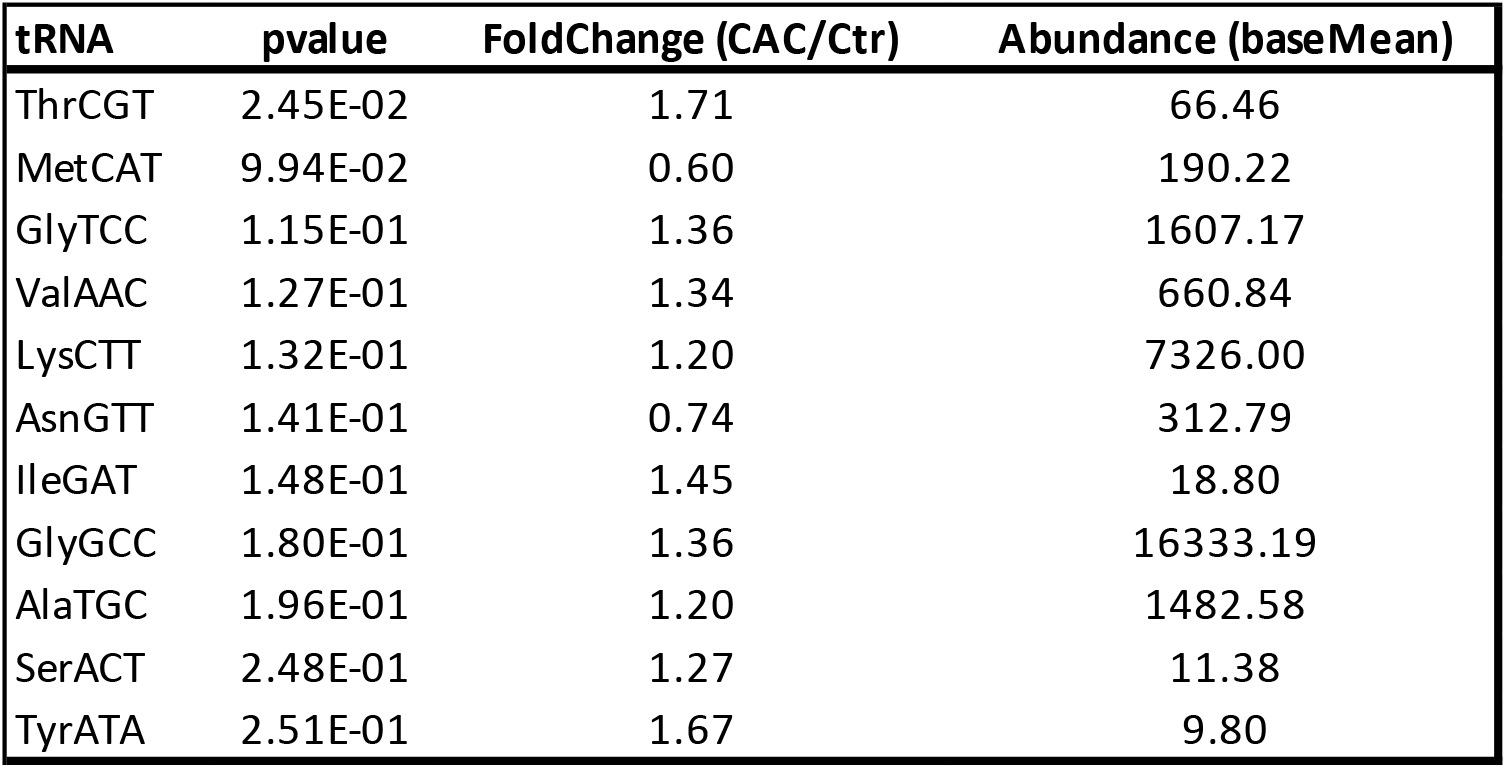
HDL-tDR changes associated with atherosclerosis, as reported by parent tRNA summary counts.

**Table S4.**
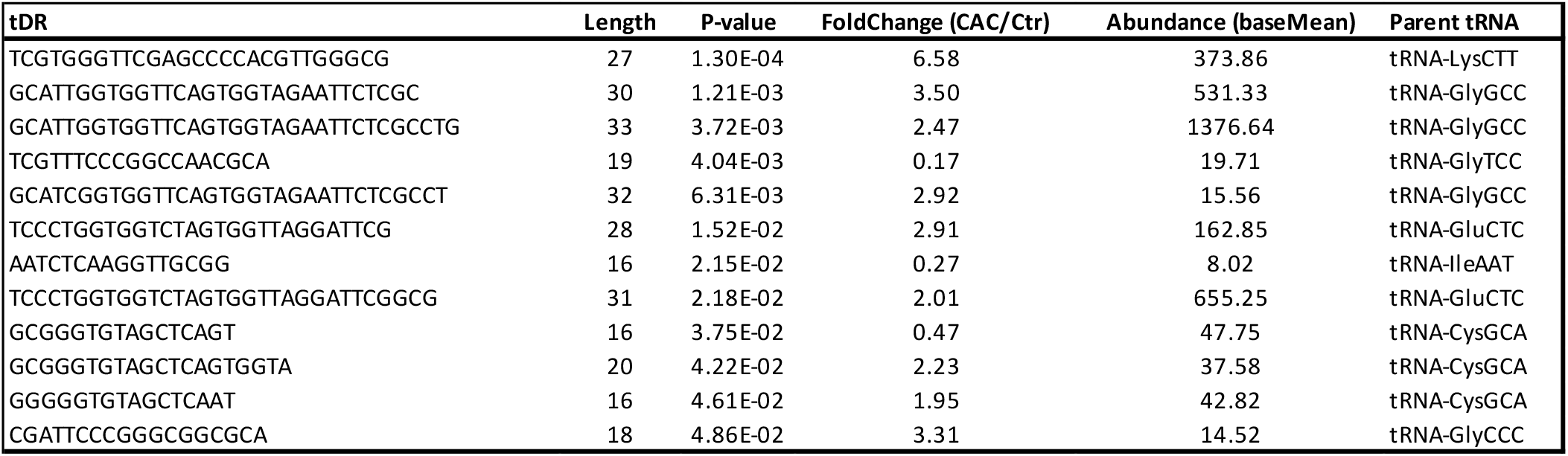
Significantly altered host HDL-tDRs in human atherosclerosis (read level).

